# Forward Simulations of Walking on Surfaces with Asymmetric Mechanical Impedance: Insights for Gait Rehabilitation

**DOI:** 10.1101/2024.10.03.616487

**Authors:** Banu Abdikadirova, Mark Price, Wouter Hoogkamer, Meghan E. Huber

**Author notes:** Banu Abdikadirova and Mark Price are co-first authors on this work. Email Addresses.

## Abstract

Gait asymmetry, prevalent in stroke survivors and various other neurological and musculoskeletal conditions, leads to abnormal joint loading, increased fall risk, and reduced walking efficiency. Traditional rehabilitation methods often fail to consistently reduce weight-bearing gait asymmetry, necessitating innovative approaches. This study explores the potential of an adjustable mechanical impedance treadmill to amplify weight-bearing asymmetries, leveraging the “error amplification” technique akin to split-belt treadmill training. We developed a 2D optimal control gait model in OpenSim to simulate walking on a rigid platform with one leg and a compliant platform, with adjustable stiffness and damping, with the other. We simulated 112 unique mechanical impedance conditions of the compliant platform and analyzed the effects of these conditions on stance time, ground reaction forces (GRFs), and muscle activations. Our results identified specific impedance parameters that can be utilized to amplify propulsion asymmetries, providing a potential new approach for gait rehabilitation post-stroke. Future work should validate these results in experimental settings and further explore optimal impedance parameters for effective gait therapy of various gait impairments.

## I. INTRODUCTION

Stroke, a leading cause of long-term disability in the U.S., induces gait impairments in 80% of cases [1]. Hemiparesis, affecting 65% of stroke survivors [2], leads to a loss of voluntary movement on the affected side, often causing asymmetric gait [3]. While gait asymmetry is often associated with stroke, it is also prevalent in neurodegenerative disorders such as Parkinson’s disease [4] and multiple sclerosis [5]. Additionally, musculoskeletal disorders and injuries such as osteoarthritis [6] and anterior cruciate ligament (ACL) injury [7] can cause gait asymmetry. Consequently, gait asymmetry is widespread across various neurological and musculoskeletal conditions.

Gait asymmetry leads to abnormal joint loading patterns [8], serves as a significant predictor of falls [9], and is associated with slower walking speeds [3] and lower gait efficiency [10], [11], potentially contributing to further adverse, longterm health outcomes such as osteoarthritis [12] and injury [13]. Moreover, the impairments caused by gait asymmetry play a role in diminishing independence when performing activities of daily life among chronic stroke survivors [14]. Therefore, addressing and reducing gait asymmetry, especially in post-stroke individuals, is a critical target for gait rehabilitation.

Conventional rehabilitation methods for minimizing gait asymmetry typically concentrate on restoring function in the affected side through resistive strength training [15] and repetitive practice of functional tasks [16]. While these approaches have shown success in enhancing patient outcomes, they exhibit inconsistency in effectively reducing gait asymmetry. In fact, despite improvements in other gait-related measures after stroke rehabilitation discharge, many individuals still experience persistent spatiotemporal asymmetry in gait [17], which tends to worsen over time [18].

Split-belt treadmill training is a promising intervention for reducing step length asymmetry. Split-belt treadmill training utilizes a dual-belt treadmill with belts moving at different speeds to amplify the existing step length asymmetry in post-stroke individuals with hemiparesis [19]. During exposure to asymmetric belt speeds, the amplified step length asymmetry gradually decreases back towards pre-exposure levels as the nervous system adapts. Upon returning to symmetric belt speeds, an aftereffect is typically observed, resulting in improved step length symmetry compared to pre-exposure. Although this aftereffect diminishes after a single bout [19], [20], improvements in step length asymmetry can persist for at least three months following structured, repetitive training [21].

Despite the promise of split-belt treadmill training, other mobility-reducing gait pathologies are resistant to this treatment, including asymmetries in temporal patterns [9] (e.g., stance vs. swing time) that can result from asymmetries in weight-bearing [5] and push-off [6] forces. The need to reduce temporal asymmetries is critical as they are most prevalent in individuals post-stroke (reports range from 48% to 82%, vs. 33% to 62% for step length asymmetry) [10], [11]. Long-term walking with weight-bearing asymmetries can lead to osteoarthritis (e.g. [12], [13]). Thus, there is a critical need for novel approaches to re-train symmetric weight-bearing behavior during gait.

We hypothesize that adjusting the dynamic behavior (e.g., mechanical impedance) of the ground could be used to amplify weight-bearing asymmetries, employing the motor learning technique used in split-belt treadmill training known as “error amplification” to induce neuromotor adaptation [22]. The recent development of treadmills capable of modulating the stiffness of the walking surface in real-time from our group and others allows for the evaluation of this hypothesis [23]–[26]. However, there remain open questions, such as which stiffness values and other mechanical impedance properties, like damping, maximize the amplification of weight-bearing asymmetries.

To address this knowledge gap, we developed an optimal control gait model to simulate walking on an adjustable mechanical impedance treadmill across a range of asymmetric stiffness and damping conditions. Human locomotion tendencies, like walking speeds and stride frequencies, can be understood as strategies to minimize metabolic cost of transport [27], [28] or effort more generally [29]. Recently, we showed that optimal control simulations aimed at minimizing muscle effort can help explain how humans adapt to split-belt treadmill walking [30]. In this paper, we used our optimal control gait model to ask what set of mechanical impedance parameters (e.g., stiffness and damping) maximally amplify weight-bearing asymmetries. Answering this question is crucial not only for designing effective experimental studies based on testable predictions, but also for developing nextgeneration variable mechanical impedance treadmills for gait rehabilitation purposes.

## II. METHODS

### A. Model Description

A modified version of a 2D locomotion model was implemented in OpenSim “2D gait.osim” [31] to calculate minimum muscle effort optimal control solutions for asymmetric ground surface stiffness. The model is 62.2 kg and 1.64 m tall, and contains 8 body segments and 10 anatomical degrees-of-freedom (DoF). The lower limb joints are actuated by 18 “DeGrooteFregly2016” Hill-type muscles [32] and the lumbar joint is actuated by an ideal coordinate actuator (Fig. 1).

**Fig 1.**
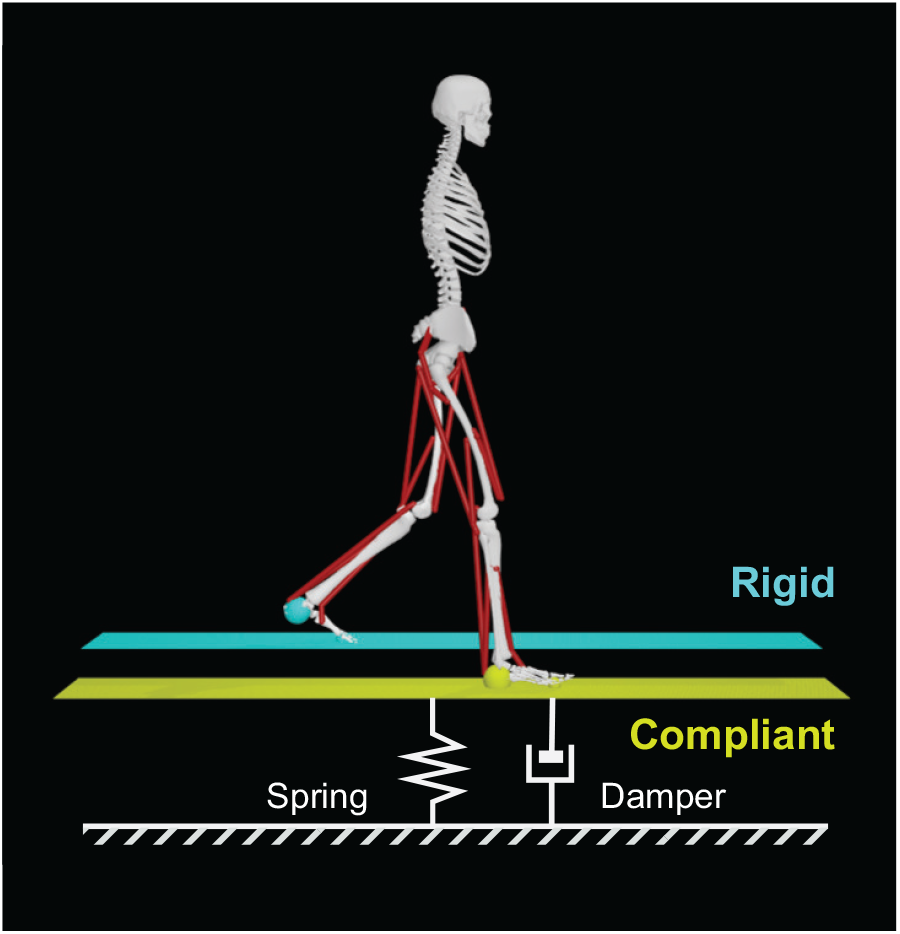
Simulation setup. The optimal control model with added platforms to simulate a dual-belt variable mechanical impedance treadmill. We simulated 112 unique impedance conditions for the compliant platform (right limb) using 16 spring and 7 damping coefficients, while the rigid platform (left limb) had fixed stiffness (1000 kN/m) and damping (1000 Ns/m) coefficients.

The model was modified by adding two platforms of negligible mass (0.1 kg) connected to the ground with linear spring-damper coordinate actuators, each with a single DoF in the vertical translation axis, to simulate the ideal behavior of the walking surface of a variable stiffness treadmill. Foot-ground contact was modeled using a smoothed Hunt-Crossley viscoelastic contact model [33] combined with a Stribeck friction model [34]. Contact spheres were located at the heel and toe of each foot with flat contact planes located on the top surface of each of the walking platforms, configured to respond only to the corresponding foot. We left this contact model with the parameters from the original model, which approximate the behavior of an athletic shoe [35], and varied the linear spring and damping coefficients of the walking platform coordinate actuators. Access to this modified model are available here (link to open-source model on simtk.org will be included here in camera-ready version of this paper).

### B. Optimal Control Simulation

The OpenSim model was simulated by solving optimal control problems with a direct collocation approach using Moco 1.0 [36]. A minimum muscle effort cost function was used, and the optimization was constrained to generate periodic strides at 1.2 m/s walking speed (within the range of the baseline muscle-reflex model configuration). The equations of motion and muscle dynamics were enforced as constraints at each of 101 nodes across the span of the gait cycle with a tolerance of 10^*−*4^.

The minimum muscle effort cost function was defined as

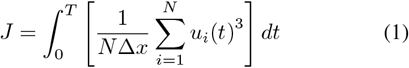

where *u*_*i*_ is the excitation of actuator *i* at time *t, N* is the total number of actuators in the model, and Δ*x* is the horizontal distance traveled by the model over one stride. This cost function approximates muscle fatigue and has been demonstrated to result in realistic kinematics, energetics, and ground reaction forces for level walking [29], [37].

Because the optimal control simulations were constrained to simulate periodic strides, transient behavior cannot be simulated; each ground mechanical impedance condition is simulated as a stable walking pattern at steady state.

### C. Simulated Mechanical Impedance Conditions

We simulated 112 unique mechanical impedance conditions for the compliant platform under the right limb, using 16 different spring and 7 damping coefficients (Fig. 2). In all simulations, the rigid platform under the left limb had fixed stiffness and damping coefficients of 1000 kN/m and 1 kNs/m, respectively.

**Fig 2.**
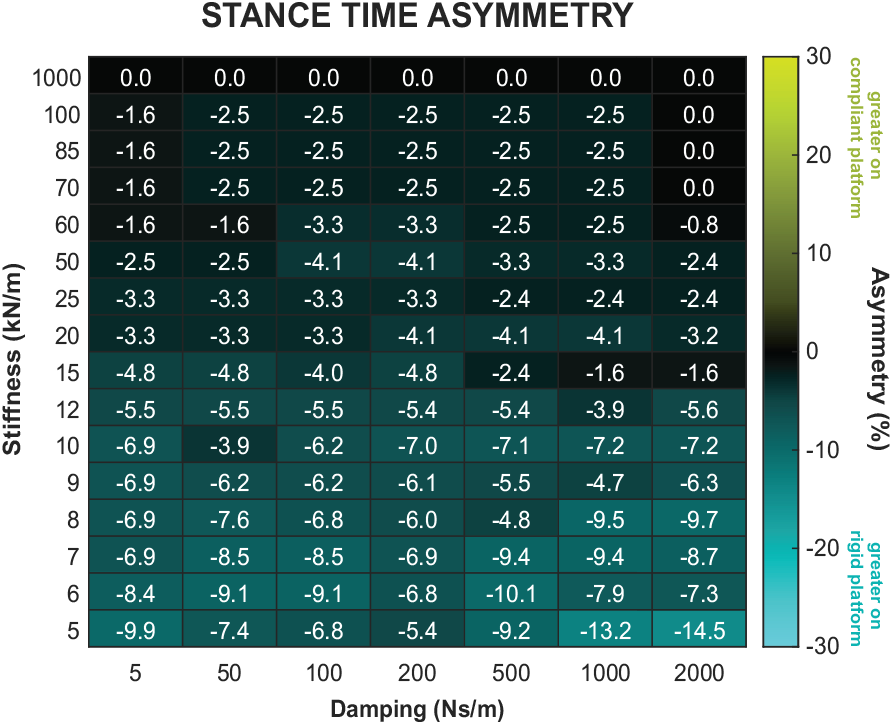
Stance Time Asymmetry. Color-coded heatmap of stance time asymmetry for all simulated mechanical impedance conditions of the compliant platform. The numbers denote the asymmetry value, and the color reflects both the magnitude (brightness) and direction (hue) of the asymmetry values as indicated by the color map.

### D. Model Analyses

Weight-bearing characteristics for each stride were quantified by measuring peak vertical, braking, and propulsive, ground reaction forces (GRFs), along with stance times for each limb. Peak vertical GRF was the maximum of the vertical GRFs, while peak propulsive and braking GRFs corresponded to the positive and negative peaks of the anteriorposterior GRFs, respectively. Stance time was defined as the duration when GRFs were non-zero.

We also measured peak levels of activation for select lower-limb muscles/muscle groups – soleus (SOL), gastrocnemius (GAS), and vasti (VAS) – that play an essential roles in forward propulsion, which is a crucial aspect of restoring symmetry in post-stroke gait rehabilitation [38]. The soleus and gastrocnemius are ankle plantarflexor muscles that contribute significantly to propulsion. The vasti muscles are knee extensors that affect leg extension during terminal stance, which also significantly influences propulsion [38].

For each simulation condition, asymmetry in the peak vertical GRFs, propulsive GRFs, braking GRFs, stance times, and muscle activations between each limb was quantified by the following ratio:

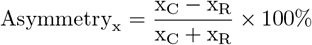

where x_C_ and x_R_ represent the measure for the compliant platform (right limb) and the rigid platform (left limb), respectively. A positive asymmetry in the measure indicates that the magnitude of the measure was higher for the right limb on the compliant platform. The opposite is true for a negative asymmetry.

## III. SIMULATION RESULTS

### A. Stance Time Asymmetry

Across all damping conditions, stance time asymmetry becomes more negative as the stiffness of the compliant platform decreases (Fig. 2). Negative stance time asymmetry indicates that the model’s stance time is longer on the rigid platform than on the compliant platform.

### B. Peak Ground Reaction Force (GRF) Asymmetries

1) *Vertical GRF Asymmetry:* Negative peak vertical GRF asymmetries were observed in simulation conditions with higher values of both stiffness and damping, while positive asymmetries generally occurred during the lower stiffness and damping conditions, except for a region with very low values (Fig. 3A).
2) *Braking GRF Asymmetry:* Positive peak braking GRF asymmetries were observed in simulation conditions with higher values of stiffness and lower values of damping. Negative asymmetries were observed in lower stiffness and higher damping conditions (Fig. 3B).
3) *Propulsive GRF Asymmetry:* Positive peak braking GRF asymmetries were observed in simulation conditions with higher values of stiffness across most damping conditions, whereas negative asymmetries were observed in lower stiffness conditions (Fig. 3C).

**Fig 3.**
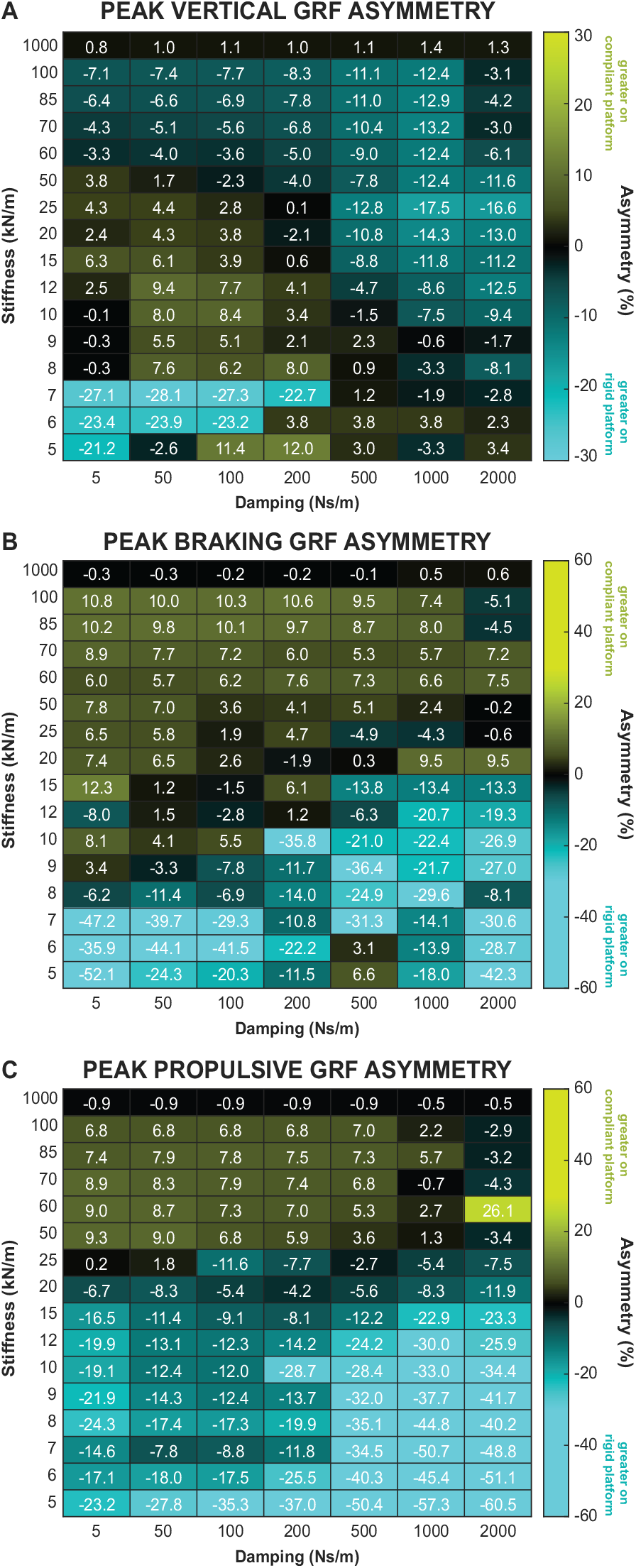
Ground reaction force (GRF) asymmetries. Color-coded heatmap of **(A)** peak vertical GRF asymmetry, **(A)** peak braking GRF asymmetry, and **(C)** peak propulsive GRF asymmetry for all simulated mechanical impedance conditions of the compliant platform. The numbers denote the asymmetry value, and the color reflects both the magnitude (brightness) and direction (hue) of the asymmetry values as indicated by the color map.

### C. Peak Muscle Activation Asymmetry

1) *Soleus (SOL) activation asymmetry:* The soleus muscle acts to plantarflex the foot about the ankle. The largest observed asymmetries in peak soleus activation were positive, indicating greater activation on the compliant platoform, and occurred in conditions with lower stiffness and damping values (Fig. 4A).
2) *Gastrocnemious (GAS) activation asymmetry:* The function of the gastrocnemius muscle group is closely related to that of the soleus muscle, as both work to plantarflex the foot. Consequently, the impact of stiffness and damping conditions on asymmetry in gastrocnemius activation mirrors that observed in the soleus muscle (Fig. 4B).
3) *Vasti (VAS) activation asymmetry:* The vasti muscle group functions to extend the knee. Positive peak asymmetries in vasti activation were observed in simulations with higher stiffness and lower damping, while negative asymmetries appeared under conditions of lower stiffness and higher damping (Fig. 4 C). This pattern mirrors the peak braking GRF asymmetry (Fig. 3B), which is consistent with the role of knee extension in braking [39].

**Fig 4.**
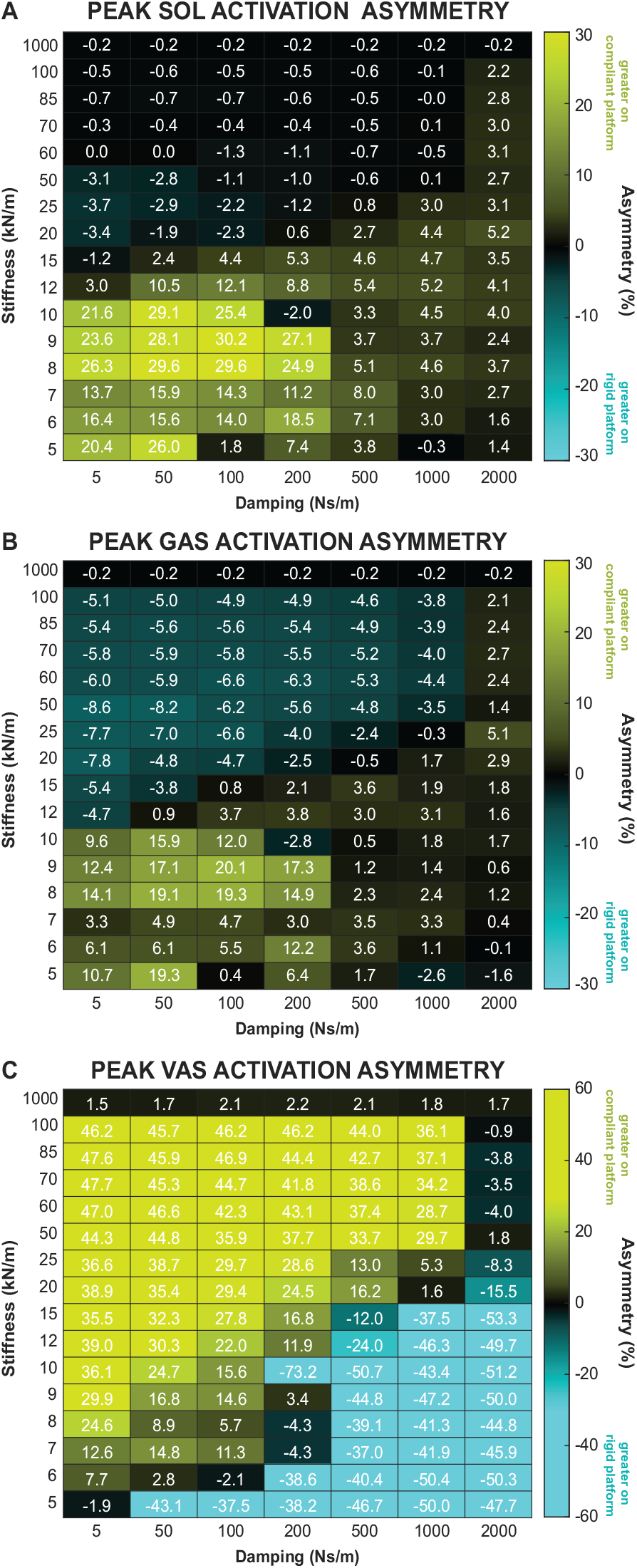
Muscle activation asymmetries. Color-coded heatmap of **(A)** peak soleus (SOL) activation asymmetry, **(A)** peak gastrocnemius (GAS) activation asymmetry, and **(C)** peak vasti (VAS) activation asymmetry for all simulated mechanical impedance conditions of the compliant platform. The numbers denote the asymmetry value, and the color reflects both the magnitude (brightness) and direction (hue) of the asymmetry values as indicated by the color map.

## IV. DISCUSSION

In this study, forward, predictive simulations of human gait demonstrate how modulating the stiffness and damping properties of a compliant platform beneath one leg, while maintaining a rigid platform under the other, affects asymmetry in relevant gait measures. By characterizing the relationship between mechanical impedance parameters and gait asymmetry, we can inform the development of new experimental protocols (e.g., approaches to amplify gait symmetry) and new hardware (e.g., variable mechanical impedance dual-belt treadmills) for gait rehabilitation purposes.

The simulation results reveal some useful guidelines for future device design. It is apparent that the useful range of stiffness and damping vary depending on the intended effect. For example, if stimulating increased plantarflexor activity is the goal of an intervention, the simulations suggest that achieving damping below 500 N-s/m is critical, regardless of how low a stiffness can be achieved by the hardware. The results also suggest that varying damping, not stiffness, may produce a wider range of behaviors for certain measures. Vertical ground reaction force asymmetry, for example, varies from +9.4 to −12.5% asymmetry across the damping range at 12 kN/m. Aside from the sharp jump in asymmetry at extremely low stiffnesses, which have been shown to be infeasible for some clinical participants to endure in existing hardware [40], this range is not achievable by changing stiffness alone.

An example of how these simulations can inform rehabilitation strategies is by identifying specific impedance parameters that amplify asymmetries in key gait measurements. For instance, when the compliant platform was set to a stiffness of 9 kN/m and damping of 50 Ns/m, the simulation produced significant asymmetry in both propulsive ground reaction forces (GRFs) and the activation of muscles contributing to propulsion (Fig. 5). Such conditions could be used to amplify existing asymmetries in stroke patients with hemiplegic gait. Since the paretic limb’s ability to generate propulsion is a critical determinant of one’s ability to walk long-distances after stroke [41], amplifying these asymmetries during training may facilitate greater neuromotor adaptation. The hypothesis is that similar to split-belt treadmill training, repeated training under these conditions could ultimately reduce gait asymmetry post-intervention, leading to improved functional outcomes for stroke survivors [21].

**Fig 5.**
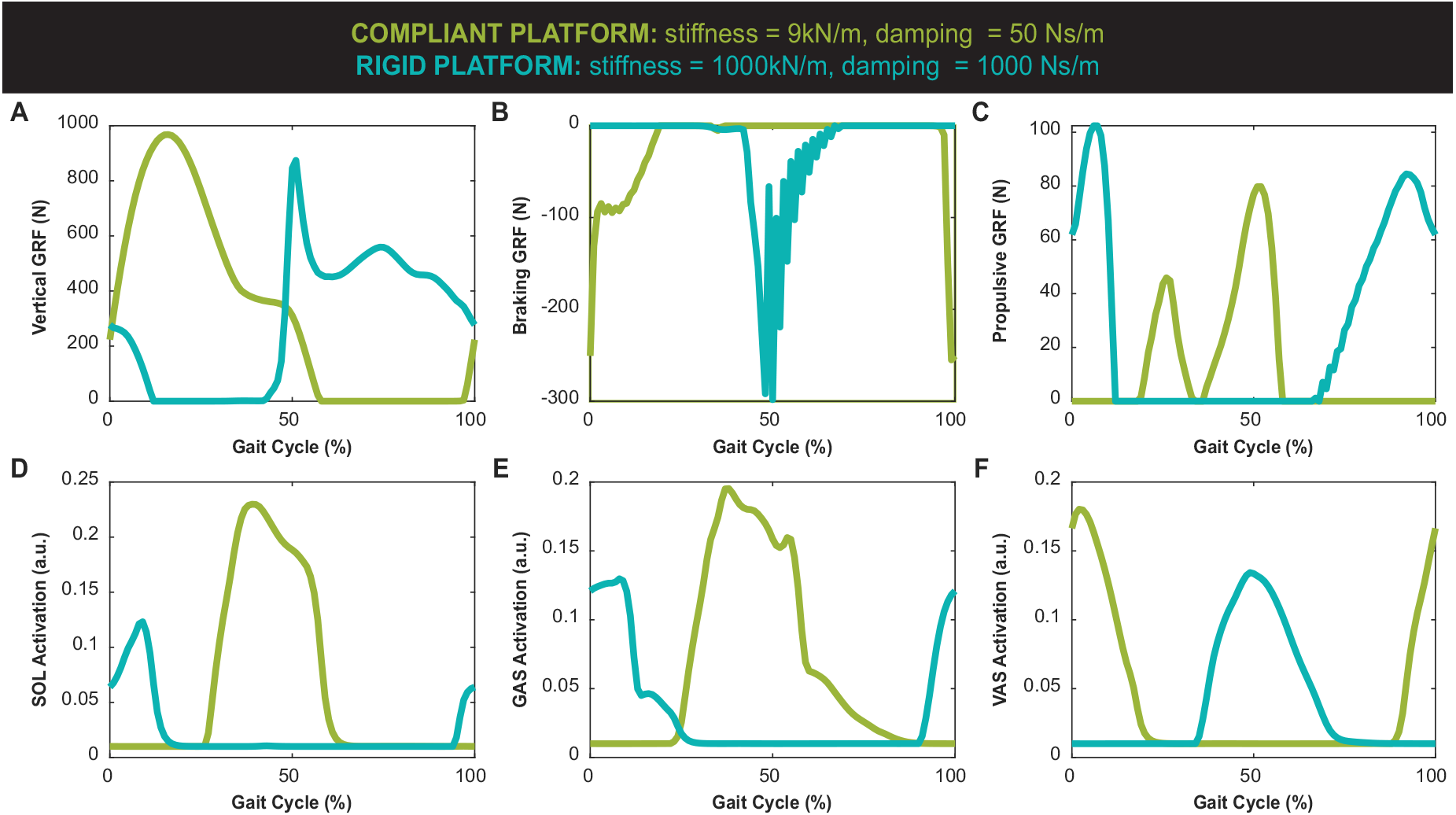
Gait measures over a gait cycle from one simulation condition that generates large asymmetries in propulsion. Profiles of **(A)** vertical GRF, **(B)** braking GRF (negative anterior-posterior GRF), **(C)** propulsive GRF (positive anterior-posterior GRF), **(D)** soleus (SOL) muscle activation, **(E)** gastrocnemius (GAS) muscle activation, and **(F)** vasti (VAS) muscle activation across one stride. The green line represents the left limb (rigid platform), while the blue line shows the right limb (compliant platform). Note that high frequency oscillations in the GRF signals occurred partly because the rigid platform was modeled with very high stiffness instead of locking the coordinate, making the force calculations highly sensitive to small deflections.

One of the significant challenges in gait rehabilitation is the heterogeneity of gait impairments, which varies not only among stroke survivors but also across individuals with different conditions causing gait asymmetries. This variability underscores the importance of characterizing the detailed landscape of gait asymmetry, as it allows for both general insights and the potential to address patient-specific asymmetries and/or coordination issues. While the use of predictive simulations to target these specific asymmetries holds promise, practical implementation in clinical settings requires further advancements in several key areas to overcome current limitations.

First, improvements in modeling are crucial. The optimal control model used in this study, though robust, has limitations. For instance, the direct collocation method does not account for gait adaptations over time, nor does it simulate the neuromotor control structures necessary for modeling changes in foot-ground contact conditions. Moreover, it does not incorporate interactions between supraspinal and peripheral neuromotor control. To address these limitations, there is a need for more advanced models capable of simulating long-term adaptations to persistent asymmetries. Techniques such as single-shooting optimal control [42], [43] and reinforcement learning [44], [45] offer potential solutions, as they can better simulate interactive perturbations and observe behavioral changes, making them suitable for modeling both the immediate and long-term effects of significant stiffness or damping perturbations.

Second, advancements in hardware development are needed. Current adjustable mechanical impedance treadmills are designed to vary stiffness but not damping (e.g., [23]– [26]). As suggested by the simulation results, the ability to independently modulate both stiffness and damping parameters could enhance the effectiveness of these devices in rehabilitation settings. Developing hardware that allows for such fine-tuned control could significantly improve the ability to address and amplify specific gait asymmetries, thereby optimizing rehabilitation outcomes for individuals with diverse gait impairments.

These two advancements – enhanced modeling and improved hardware – will mutually benefit from each other. The introduction of variable impedance treadmills will allow us to conduct novel experiments that assess how prolonged exposure to walking on surfaces with asymmetric stiffness and damping properties influences muscle activity, as well as lower limb kinematics and kinetics. Such experiments will enable us to better understand the long-term adaptation processes, as well as validate and refine the models based on real-world data.

## REFERENCES

[1] P. W. Duncan, R. Zorowitz, B. Bates, J. Y. Choi, J. J. Glasberg, G. D. Graham, R. C. Katz, K. Lamberty, and D. Reker, “Management of adult stroke rehabilitation care: a clinical practice guideline,” Stroke, vol. 36, no. 9, pp. e100–e143, 2005.

[2] J. H. Cauraugh and S. B. Kim, “Chronic stroke motor recovery: duration of active neuromuscular stimulation,” Journal of the Neurological Sciences, vol. 215, no. 1-2, pp. 13–19, 2003.

[3] K. K. Patterson, I. Parafianowicz, C. J. Danells, V. Closson, M. C. Verrier, W. R. Staines, S. E. Black, and W. E. McIlroy, “Gait asymmetry in community-ambulating stroke survivors,” Archives of Physical Medicine and Rehabilitation, vol. 89, no. 2, pp. 304–310, 2008.

[4] B. W. Fling, C. Curtze, and F. B. Horak, “Gait asymmetry in people with parkinson’s disease is linked to reduced integrity of callosal sensorimotor regions,” Frontiers in Neurology, vol. 9, p. 215, 2018.

[5] M. Plotnik, J. M. Wagner, G. Adusumilli, A. Gottlieb, and R. T. Naismith, “Gait asymmetry, and bilateral coordination of gait during a six-minute walk test in persons with multiple sclerosis,” Scientific reports, vol. 10, no. 1, p. 12382, 2020.

[6] G. J. Farkas, B. R. Schlink, L. F. Fogg, K. C. Foucher, M. A. Wimmer, and N. Shakoor, “Gait asymmetries in unilateral symptomatic hip osteoarthritis and their association with radiographic severity and pain,” Hip International, vol. 29, no. 2, pp. 209–214, 2019.

[7] S. A. Garcia, S. R. Brown, M. Koje, C. Krishnan, and R. M. Palmieri-Smith, “Gait asymmetries are exacerbated at faster walking speeds in individuals with acute anterior cruciate ligament reconstruction,” Journal of Orthopaedic Research, vol. 40, no. 1, pp. 219–230, 2022.

[8] S. Marrocco, L. D. Crosby, I. C. Jones, R. F. Moyer, T. B. Birmingham, and K. K. Patterson, “Knee loading patterns of the non-paretic and paretic legs during post-stroke gait,” Gait & posture, vol. 49, pp. 297– 302, 2016.

[9] K. Bower, S. Thilarajah, Y.-H. Pua, G. Williams, D. Tan, B. Mentiplay, L. Denehy, and R. Clark, “Dynamic balance and instrumented gait variables are independent predictors of falls following stroke,” Journal of Neuroengineering and Rehabilitation, vol. 16, pp. 1–9, 2019.

[10] N. Sànchez and J. M. Finley, “Individual differences in locomotor function predict the capacity to reduce asymmetry and modify the energetic cost of walking poststroke,” Neurorehabilitation and Neural Repair, vol. 32, no. 8, pp. 701–713, 2018.

[11] G. Balbinot, C. P. Schuch, H. Bianchi Oliveira, and L.A. Peyré-Tartaruga, “Mechanical and energetic determinants of impaired gait following stroke: segmental work and pendular energy transduction during treadmill walking,” Biology Open, vol. 9, no. 7, p. bio051581, 2020.

[12] W. Li, Y. Li, Q. Gao, J. Liu, Q. Wen, S. Jia, F. Tang, L. Mo, Y. Zhang, M. Zhai et al., “Change in knee cartilage components in stroke patients with genu recurvatum analysed by zero te mr imaging,” Scientific reports, vol. 12, no. 1, p. 3751, 2022.

[13] J. E. Harris, J. J. Eng, D. S. Marigold, C. D. Tokuno, and C. L. Louis, “Relationship of balance and mobility to fall incidence in people with chronic stroke,” Physical Therapy, vol. 85, no. 2, pp. 150–158, 2005.

[14] A. Schmid, P. W. Duncan, S. Studenski, S. M. Lai, L. Richards, S. Perera, and S. S. Wu, “Improvements in speed-based gait classifications are meaningful,” Stroke, vol. 38, no. 7, pp. 2096–2100, 2007.

[15] L. Glasser, “Effects of isokinetic training on the rate of movement during ambulation in hemiparetic patients,” Physical Therapy, vol. 66, no. 5, pp. 673–676, 1986.

[16] D. A. Gelber, B. Josefczyk, D. Herrman, D. C. Good, and S. J. Verhulst, “Comparison of two therapy approaches in the rehabilitation of the pure motor hemiparetic stroke patient,” Journal of Neurologic Rehabilitation, vol. 9, no. 4, pp. 191–196, 1995.

[17] G. M. Rozanski, J. S. Wong, E. L. Inness, K. K. Patterson, and A. Mansfield, “Longitudinal change in spatiotemporal gait symmetry after discharge from inpatient stroke rehabilitation,” Disability and Rehabilitation, vol. 42, no. 5, pp. 705–711, 2020.

[18] G. Turnbull and J. Wall, “Long-term changes in hemiplegic gait,” Gait & Posture, vol. 3, no. 4, pp. 258–261, 1995.

[19] D. S. Reisman, R. Wityk, K. Silver, and A. J. Bastian, “Locomotor adaptation on a split-belt treadmill can improve walking symmetry post-stroke,” Brain, vol. 130, no. 7, pp. 1861–1872, 2007.

[20] D. S. Reisman, R. Wityk, K. Silver, and A. J. Bastian, “Split-belt treadmill adaptation transfers to overground walking in persons poststroke,” Neurorehabilitation and Neural Repair, vol. 23, no. 7, pp. 735–744, 2009.

[21] D. S. Reisman, H. McLean, J. Keller, K. A. Danks, and A. J. Bastian, “Repeated split-belt treadmill training improves poststroke step length asymmetry,” Neurorehabilitation and Neural Repair, vol. 27, no. 5, pp. 460–468, 2013.

[22] D. J. Reinkensmeyer and J. L. Patton, “Can robots help the learning of skilled actions?” Exercise and Sport Sciences Reviews, vol. 37, no. 1, pp. 43–51, 2009.

[23] J. Skidmore, A. Barkan, and P. Artemiadis, “Variable stiffness treadmill (VST): System development, characterization, and preliminary experiments,” IEEE/ASME Transactions on Mechatronics, vol. 20, no. 4, pp. 1717–1724, 2015.

[24] E. Hernandez, C. Warhmund, K. Lamoureux, E. Lee, I. Sanchez, W. Matthews, and A. Jafari, “A novel treadmill that can bilaterally adjust the vertical surface stiffness,” IEEE/ASME Transactions on Mechatronics, vol. 23, no. 5, pp. 2338–2346, 2018.

[25] V. Chambers, B. Hobbs, W. Gaither, A. Zhou, C. Karakasis, P. Artemiadis et al., “The variable stiffness treadmill (vst) 2: Development and validation of a unique tool to investigate locomotion on compliant terrains,” Journal of Mechanisms and Robotics, pp. 1–11, 2024.

[26] M. Price, D. Locurto, B. Abdikadirova, M. E. Huber, and W. Hoogkamer, “Adjusst: An adjustable surface stiffness treadmill,” bioRxiv, pp. 2024–03, 2024.

[27] H. J. Ralston, “Energetics of human walking,” in Neural Control of Locomotion, R. M. Herman, S. Grillner, P. S. G. Stein, and S. D. G, Eds. Boston, MA: Springer, 1976, pp. 77–98.

[28] Z. M. Y. and R. C. W., “Predicting metabolic cost of level walking,” European Journal of Applied Physiology and Occupational Physiology, vol. 38, pp. 215–223, 1978.

[29] M. Ackermann and A. J. van den Bogert, “Optimality principles for model-based prediction of human gait,” Journal of Biomechanics, vol. 43, no. 6, pp. 1055–1060, 2010.

[30] M. Price, M. E. Huber, and W. Hoogkamer, “Minimum effort simulations of split-belt treadmill walking exploit asymmetry to reduce metabolic energy expenditure,” Journal of Neurophysiology, vol. 129, no. 4, pp. 900–913, 2023.

[31] S. L. Delp, F. C. Anderson, A. S. Arnold, P. Loan, A. Habib, C. T. John, E. Guendelman, and D. G. Thelen, “Opensim: Open-source software to create and analyze dynamic simulations of movement,” IEEE Transactions on Biomedical Engineering, vol. 54, no. 11, 2007.

[32] F. De Groote, A. L. Kinney, A. V. Rao, and B. J. Fregly, “Evaluation of direct collocation optimal control problem formulations for solving the muscle redundancy problem,” Annals of Biomedical Engineering, vol. 44, pp. 2922–2936, 2016.

[33] G. Serrancolí, A. Falisse, C. Dembia, J. Vantilt, K. Tanghe, D. Lefeber, I. Jonkers, J. De Schutter, and F. De Groote, “Subject-exoskeleton contact model calibration leads to accurate interaction force predictions,” IEEE Transactions on Neural Systems and Rehabilitation Engineering, vol. 27, no. 8, pp. 1597–1605, 2019.

[34] M. A. Sherman, A. Seth, and S. L. Delp, “Simbody: Multibody dynamics for biomedical research,” Procedia IUTAM, vol. 2, pp. 241– 261, 2011.

[35] P. Aerts and D. D. Clercq, “Deformation characteristics of the heel region of the shod foot during a simulated heel strike: The effect of varying midsole hardness,” Journal of Sports Sciences, vol. 11, no. 5, pp. 449–461, 1993.

[36] C. L. Dembia, N. A. Bianco, A. Falisse, J. L. Hicks, and S. L. Delp, “Opensim moco: Musculoskeletal optimal control,” PLoS Computational Biology, vol. 16, no. 12, p. e1008493, 2020.

[37] R. H. Miller, B. R. Umberger, J. Hamill, and G. E. Caldwell, “Evaluation of the minimum energy hypothesis and other potential optimality criteria for human running,” Proceedings of the Royal Society B: Biological Sciences, vol. 279, no. 1733, pp. 1498–1505, 2012.

[38] S. A. Roelker, M. G. Bowden, S. A. Kautz, and R. R. Neptune, “Paretic propulsion as a measure of walking performance and functional motor recovery post-stroke: A review,” Gait & Posture, vol. 68, pp. 6–14, 2019.

[39] R. G. Ellis, B. J. Sumner, and R. Kram, “Muscle contributions to propulsion and braking during walking and running: insight from external force perturbations,” Gait & Posture, vol. 40, no. 4, pp. 594– 599, 2014.

[40] V. Chambers and P. Artemiadis, “Using robot-assisted stiffness perturbations to evoke aftereffects useful to post-stroke gait rehabilitation,” Frontiers in Robotics and AI, vol. 9, p. 1073746, 2023.

[41] L. N. Awad, S. A. Binder-Macleod, R. T. Pohlig, and D. S. Reisman, “Paretic propulsion and trailing limb angle are key determinants of long-distance walking function after stroke,” Neurorehabilitation and Neural Repair, vol. 29, no. 6, pp. 499–508, 2015.

[42] A. D. Russo, D. Stanev, S. Armand, and A. Ijspeert, “Sensory modulation of gait characteristics in human locomotion: A neuromusculoskeletal modeling study,” PLoS Computational Biology, vol. 17, no. 5, p. e1008594, 2021.

[43] C. F. Ong, T. Geijtenbeek, J. L. Hicks, and S. L. Delp, “Predicting gait adaptations due to ankle plantarflexor muscle weakness and contracture using physics-based musculoskeletal simulations,” PLoS computational biology, vol. 15, no. 10, p. e1006993, 2019.

[44] N. Seethapathi, B. Clark, and M. Srinivasan, “Exploration-based learning of a step to step controller predicts locomotor adaptation,” bioRxiv, March 2021.

[45] L. de Vree and R. Carloni, “Deep reinforcement learning for physics-based musculoskeletal simulations of healthy subjects and transfemoral prostheses’ users during normal walking,” IEEE Transactions on Neural Systems and Rehabilitation Engineering, vol. 29, pp. 607– 618, 2021.

